# Bayesian Optimized sample-specific Networks Obtained By Omics data (BONOBO)

**DOI:** 10.1101/2023.11.16.567119

**Authors:** Enakshi Saha, Viola Fanfani, Panagiotis Mandros, Marouen Ben-Guebila, Jonas Fischer, Katherine Hoff-Shutta, Kimberly Glass, Dawn Lisa DeMeo, Camila Lopes-Ramos, John Quackenbush

**Affiliations:** Department of Biostatistics, Harvard T.H. Chan School of Public Health, Boston, MA; Channing Division of Network Medicine, Brigham and Women’s Hospital, Boston, MA, USA; Department of Medicine, Harvard Medical School, Boston, MA, USA; Department of Data Science, Dana-Farber Cancer Institute, Boston, MA, USA

**Keywords:** Gene regulatory network, Co-expression, individual-specific network, Bayesian inference, posterior distribution

## Abstract

Gene regulatory networks (GRNs) are effective tools for inferring complex interactions between molecules that regulate biological processes and hence can provide insights into drivers of biological systems. Inferring co-expression networks is a critical element of GRN inference as the correlation between expression patterns may indicate that genes are coregulated by common factors. However, methods that estimate co-expression networks generally derive an aggregate network representing the mean regulatory properties of the population and so fail to fully capture population heterogeneity. To address these concerns, we introduce BONOBO (Bayesian Optimized Networks Obtained By assimilating Omics data), a scalable Bayesian model for deriving individual sample-specific co-expression networks by recognizing variations in molecular interactions across individuals. For every sample, BONOBO assumes a Gaussian distribution on the log-transformed centered gene expression and a conjugate prior distribution on the sample-specific co-expression matrix constructed from all other samples in the data. Combining the sample-specific gene expression with the prior distribution, BONOBO yields a closed-form solution for the posterior distribution of the sample-specific co-expression matrices, thus making the method extremely scalable. We demonstrate the utility of BONOBO in several contexts, including analyzing gene regulation in yeast transcription factor knockout studies, prognostic significance of miRNA-mRNA interaction in human breast cancer subtypes, and sex differences in gene regulation within human thyroid tissue. We find that BONOBO outperforms other sample-specific co-expression network inference methods and provides insight into individual differences in the drivers of biological processes.

**Graphical Abstract:** 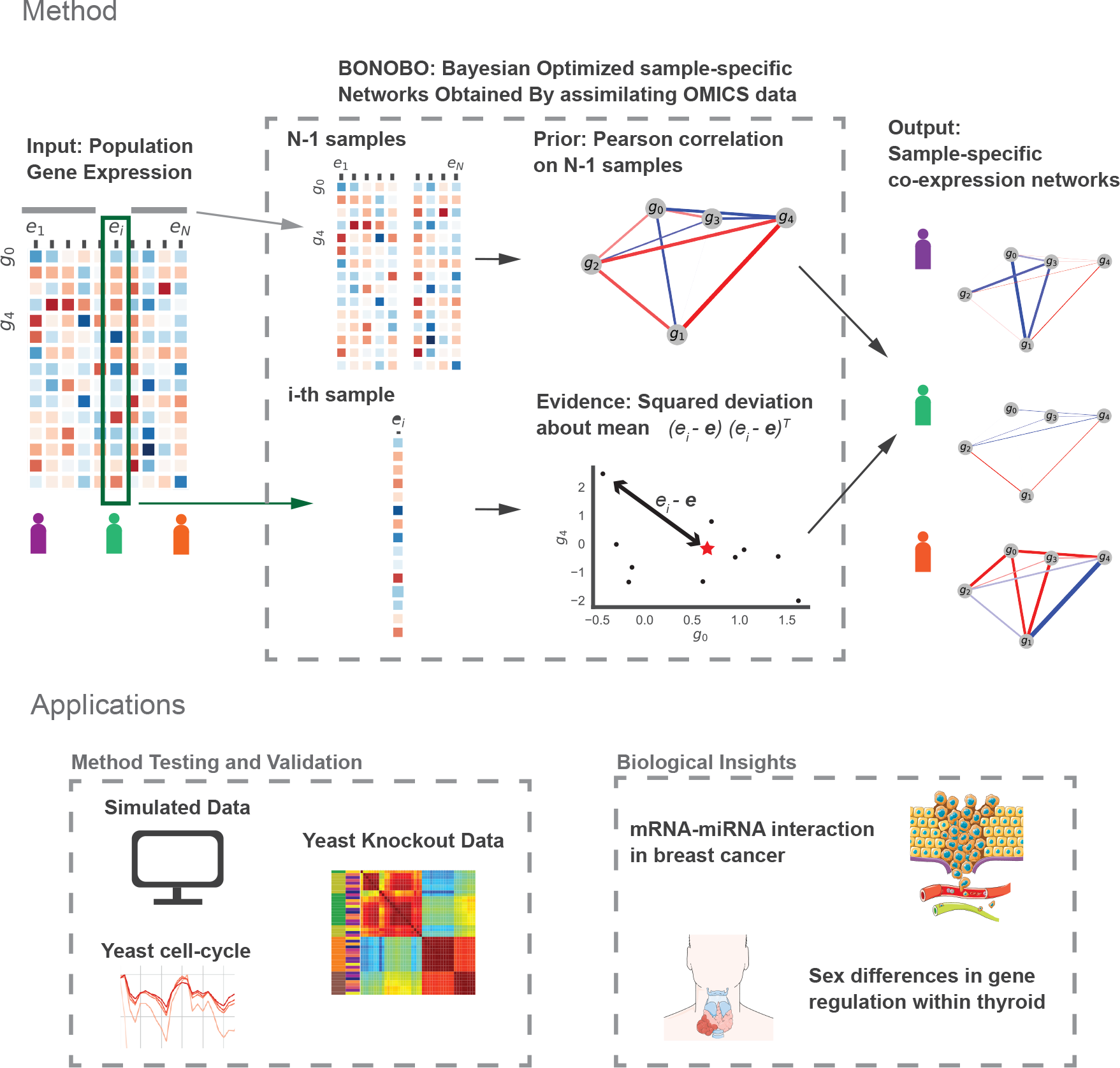

## 1 Introduction

The majority of human traits and diseases are driven not by individual genes, but by networks of genes and proteins interacting with each other [1]. Understanding how genes interact and cooperate under different conditions is a central challenge in deciphering the complexities of cellular processes and their dysregulation in various diseases. While differential expression analysis with conventional tools such as “limma” (or “voom”)[2] enables us to adjust for the effects of these covariates, differences found in transcription levels alone often fail to explain biological differences between the cohorts being compared [3].

Co-expression networks, which represent the coordinated expression patterns of genes across diverse biological samples, can provide insights into processes that are simultaneously activated in different biological states. However, most methods for constructing co-expression networks estimate an aggregate network for the entire population [4, 5, 6], thus failing to capture the heterogeneous, context-specific gene interactions present within individual samples. Trying to overcome these limitations, methods to infer sample-specific co-expression networks have been proposed, such as Single Pearson Correlation Coefficient (SPCC) [7, 8] and Linear Interpolation to Obtain Network Estimates for Single Samples (LIONESS) [9]. However, these methods produce co-expression matrices that are not positive definite and/or where the estimated correlation values are assigned outside the defined range for correlation measures (for example, [-1,1] for Pearson’s correlation coefficient). This non-positive definiteness can pose significant challenges in downstream analyses, as it violates the fundamental assumptions of correlation networks and can lead to misleading interpretations. Alternatively, other methods designed for personalized characterization of diseases through sample-specific networks [10] and cancer-specific or group-specific networks [11] represent differential networks with respect to an external reference population and hence are susceptible to varying inference depending on the reference sample used.

We develop BONOBO (Bayesian Optimized Networks Obtained By assimilating Omics data), an empirical Bayesian model that derives individual sample-specific co-expression networks (Figure 1), thus facilitating the discovery of differentially co-regulated gene pairs between different conditions and/or phenotypes, while eliminating the effects of confounders. BONOBO derives positive definite co-expression networks from input data alone, without using any external reference datasets. This distinctive feature enables BONOBO to capture correlation structures that remain consistent and comparable across diverse datasets and multiple batches, providing a robust tool for network analysis. BONOBO derives a posterior probability distribution for individual correlation matrices, allowing us to test the hypothesis of whether any two pairs of genes have a non-zero correlation, within an individual sample in the data. Based on the results of these hypotheses-testing we can infer individual sample-specific sparse co-expression networks by pruning out non-significant edges. This is particularly important for interpretability as empirical data suggests that biological gene networks are sparsely connected[12].

**Figure 1:**
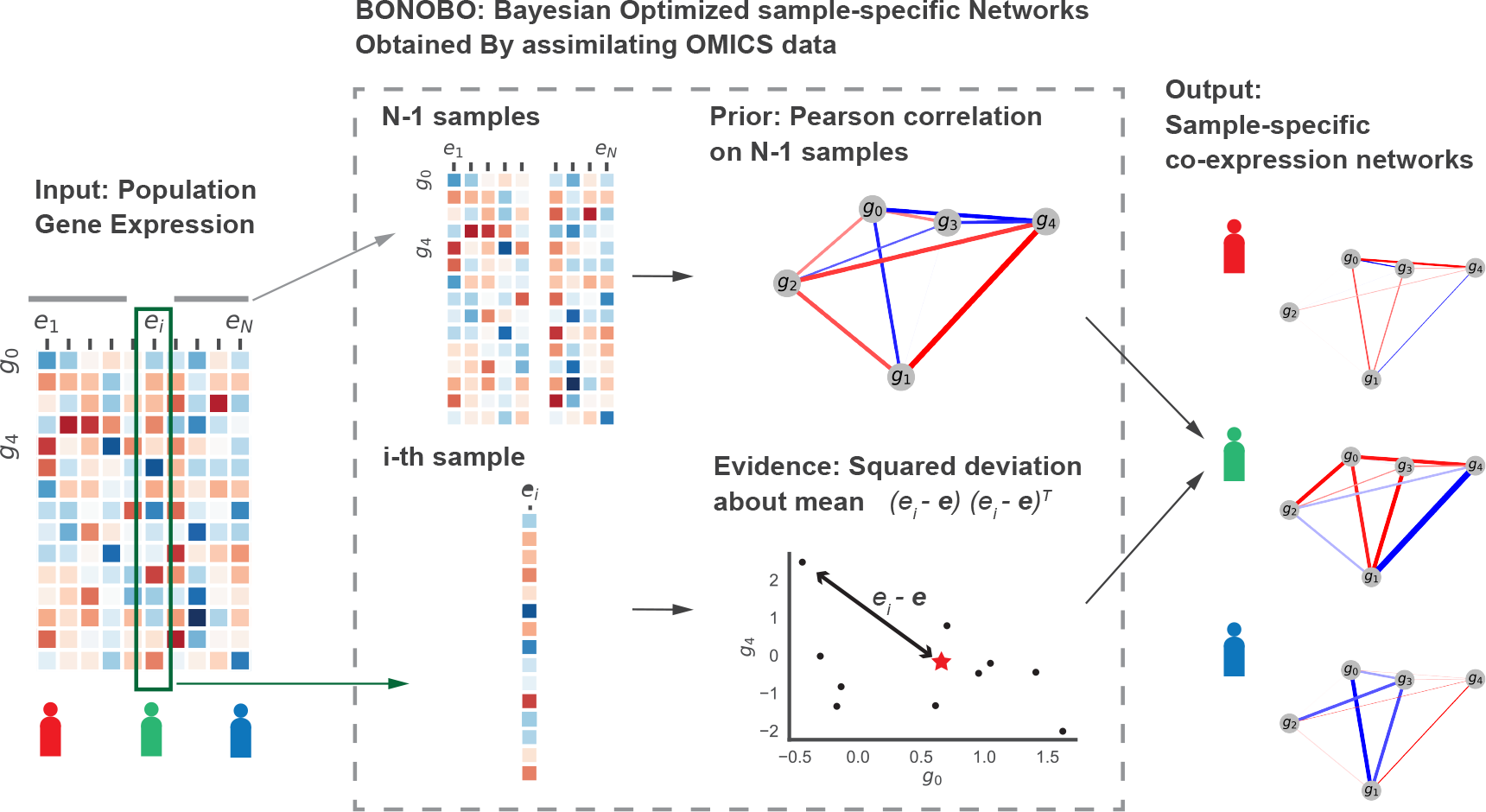
Schematic diagram of BONOBO: BONOBO requires a gene expression matrix as input, from which we would like to extract sample-specific correlation networks. Then, for each of the samples, BONOBO infers the network by using both the Pearson correlation matrix computed on *N* − 1 samples and the sample specific squared-deviation about the mean. BONOBO outputs *N* co-expression networks, one for each sample, and the associated p-values for each of the gene-gene estimated edges.

One of the key strengths of BONOBO lies in its ability to capture the inherent heterogeneity in co-expression patterns among individuals within a population, that can be attributed to a range of biological and environmental elements. For instance, when comparing aggregate co-expression networks between conditions, such as distinguishing between health and disease or male and female samples, results are frequently confounded by the population’s heterogeneity stemming from nuisance parameters, such as batch effects and/or confounding clinical covariates such as race and age. BONOBO’s individual sample-specific approach explicitly models this heterogeneity, enabling a deeper understanding of the gene networks underlying distinct biological states. In addition, individual sample-specific co-expression networks derived by BONOBO can also be used as inputs to methods for inferring gene regulatory networks that require a correlation matrix as input, such as PANDA (Passing Messages between Biological Networks to Refine Predicted Interactions) [3], OTTER (Optimize To Estimate Regulation) [13] and EGRET (Estimating the Genetic Regulatory Effect on Transcription factors) [14] to infer sample-specific gene regulatory networks. These methods infer bipartite gene regulatory networks consisting of directed regulatory edges from regulators such as transcription factors (TF) to their target genes, by combining gene co-expression matrices with individual-specific TF-motif and/or chromatin accessibility data.

We demonstrate the advantages of BONOBO using several simulated and real datasets. First, we used simulated data to compare BONOBO’s performance with other state-of-the-art methods. We then used pseudo-bulk gene expression data from knockout experiments in yeast cells and show that BONOBO captures global properties of each yeast strain and also distinguishes the sample-specific effects of single transcription factor knockouts. Next, we examined the interaction between miRNA and mRNA expression in various human breast cancer subtypes using individual-specific co-expression networks derived by BONOBO and find that the correlation patterns between miRNA expression and immune pathways have prognostic significance in luminal A and luminal B breast cancer subtypes. In a final application, we study sex differences in gene regulation within human thyroid. Empirical data indicate that females are three times more likely to develop some types of thyroid conditions over their lifetime than males [15]. Using BONOBO networks as inputs to PANDA, we infer individual-specific gene regulatory networks and compare these between males and females, identifying regulatory differences in immune response, cell proliferation, and metabolic processes, thereby providing a possible mechanism for sex bias in incidence rates of various thyroid conditions such as hypothyroidism and Hashimoto’s disease.

BONOBO is available as open-source code in Python through the Network Zoo package (netZooPy v0.9.17; netzoo.github.io) [16].

## 2 Methods

### 2.1 BONOBO

Let *x*_1_, *x*_2_, …, *x*_*N*_ ∼ ℝ^*g*^ denote the log-transformed bulk gene expression values of *g* genes for *N* samples. Let us assume that for every sample *i* ∈ {1, 2, …, *N*}, the centered expression vector 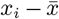 follows a multivariate normal distribution with mean zero and an unknown sample-specific covariance matrix *V*_*i*_,

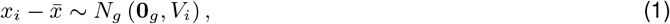

where **0**_*g*_ ∈ ℝ^*g*^ denotes a vector of all zeros and 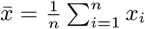 denotes the mean expression across all samples. Our objective is to estimate *V*_*i*_, the sample-specific covariance matrix of gene expression for the *i*-th sample, ∀*i*.

We assume that for every sample *i* ∈ {1, 2, …, *N* }, the sample-specific covariance matrix *V*_*i*_ follows an inverse Wishart prior distribution, given all other samples *j* ∈ {1, 2, …, *N* } \ {*i*},

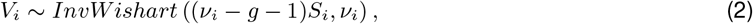

where *v*_*i*_ ≥ *g* + 1 denotes the degree of freedom and *S*_*i*_ denotes the sample covariance matrix computed from *N* − 1 samples excluding the *i*-th sample. Under this assumption, the prior mean of the covariance matrix for the *i*-th sample is 𝔼[*V*_*i*_] = *S*_*i*_. In other words, we assume that the correlation between any pair of genes for each individual is similar to the correlation between these same pair of genes across the entire population on average.

The inverse Wishart distribution is a conjugate prior for the covariance matrix of a multivariate normal distribution. Therefore, under the above prior specification, the posterior distribution of the sample-specific covariance matrix *V*_*i*_ also turns out to be an inverse Wishart distribution, as described by the following theorem.

#### Theorem 1

*Under assumptions* (1) *and* (2), *the posterior distribution of V*_*i*_ *is*

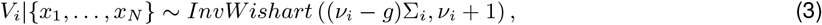

*where* 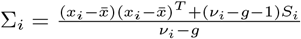 *denotes the posterior mean of V*_*i*_.

The proof of the above theorem is given in the appendix S1.1.1.

From (3) we observe that the posterior mean of *V*_*i*_, the covariance matrix of the *i*-th sample is a linear combination of the prior mean *S*_*i*_, which summarizes information from all other samples excluding the *i*-th sample and a sample-specific component 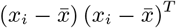, which summarizes the association between pairwise genes within the *i*-th sample alone:

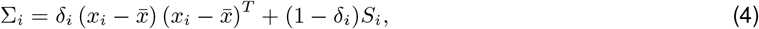

where 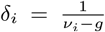. Since *v*_*i*_− *g* ≥ 1, we have 0 ≤*δ*_*i*_ ≤ 1, which represents the relative contributions of the sample-specific information and the prior information, while estimating the posterior mean of *V*_*i*_.

As sample size *n* increases, the strong law of large numbers implies 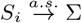, where σ denotes the population covariance matrix. Thus the hyperparameter *δ*_*i*_ quantifies the contribution of the sample-specific information in the posterior mean, while the complement 1 − *δ*_*i*_ quantifies the contribution of the population covariance matrix σ. For homogeneous populations we recommend using a smaller value of *δ*_*i*_, or equivalently, a larger value of 1 − *δ*_*i*_, as this would increase the contribution of the population covariance σ and give robust estimates of the sample-specific covariance *V*_*i*_. On the other hand, if the *i*-th sample is an outlier with respect to the rest of the population, we recommend using a large value of *δ*_*i*_, thereby decreasing estimation bias. Alternatively, we can set *δ*_*i*_ = *δ*, ∀*i* to some arbitrary value between (0, 1). In the following section we describe a computationally inexpensive data-driven empirical procedure for calibrating *δ*_*i*_ for every sample.

#### 2.1.1 Fixing Prior Degrees of Freedom

The hyperparameter 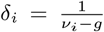 is a one-to-one function of the prior degrees of freedom *v*_*i*_. Hence in order to calibrate *δ*_*i*_, it suffices to estimate *v*_*i*_ for every sample *i* in the data. The following lemma provides a data-driven approach for calibrating *v*_*i*_.

##### Lemma 1

*Under assumption* (2), *prior variance of the k-th diagonal entry of V*_*i*_ *(denoted by* 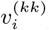*) would be*

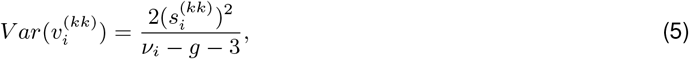

*where* 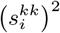 *denotes the k-th diagonal entry of S*_*i*_.

The above lemma is a direct consequence of the properties of inverse Wishart distribution [17].

From (5), summing over *k* = 1, …, *g*, i.e., over all genes, we get

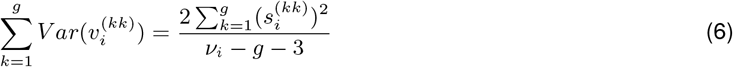

Simplifying the above equation gives us,

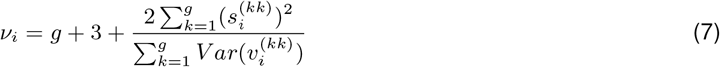

For every sample *i*, the right side of (7) is known except for 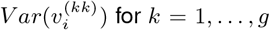 for *k* = 1, …, *g*. We can approximate this value from the data as follows:

1. Get *n* estimates of the variance of the *k*-th gene by leaving out one sample at a time: 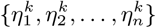, where 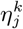 denotes the variance of *k*-th coordinates of {*x*_1_, …, *x*_*N*_ } \ *x*_*j*_ .
2. Estimate 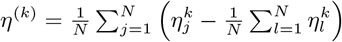, the variance of 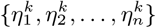.

Replacing 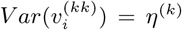 on the right hand side of equation (7) gives us a data-driven estimate of the prior degrees of freedom *v*_*i*_. Thus the estimate of the hyperparameter *δ*_*i*_ becomes

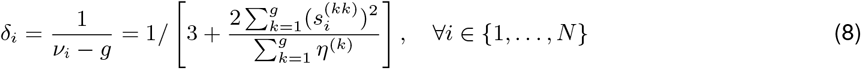

In section S1.2.5, we illustrate, via simulation experiments, that this data-driven empirical approach of calibrating *δ*_*i*_ delivers performance on par with the optimal performance achieved by fixing *δ*_*i*_ = *δ*, ∀*i* to any arbitrary value.

#### 2.1.2 Hypothesis Testing

For every sample *i* we can derive a 100(1 − *α*)% posterior credible region for the correlation between any pair of genes as follows: first we compute the posterior variance of the covariance between any two pair of genes using the following lemma, which is a direct consequence of the properties of inverse Wishart distribution [17]. For simplicity we remove the sample index *i*.

##### Lemma 2

*Let v*_*jk*_ *denote the covariance between the j-th and the k-th gene. Under assumptions* (1) *and* (2), *the posterior variance of v*_*jk*_ *would be*

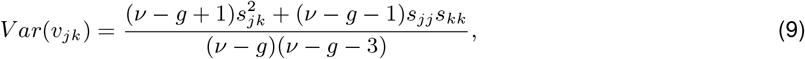

*where s*_*kk*_ *denotes the k-th diagonal entry of the prior mean S and s*_*jk*_ *denotes the* (*j, k*)*-th off-diagonal entry (corresponding to the j-th row and the k-th column) of S*.

Using the above lemma we can compute an approximate 100(1 − *α*)% posterior credible region for *v*_*jk*_ as (*σ*_*jk*_ − *ψ*_*jk*_*z*_(1*−α/*2)_, *σ*_*jk*_ + *ψ*_*jk*_*z*_(1*−α/*2)_), where 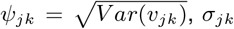 is the posterior mean of *v*_*jk*_ and *z*_(1*−α/*2)_ denotes the (1−*α/*2)-th quantile of the standard normal distribution.

Equivalently, for every pair of genes (*j, k*), we can reject the null hypothesis *H*_0_ : *v*_*jk*_ = 0, in favor of the alternative hypothesis *H*_1_ : *v*_*jk*_ ≠ 0 at significance level *α*, provided 2 (1 − Φ(*σ*_*jk*_*/ψ*_*jk*_))≤ *α*, where Φ denotes the cumulative distribution function of the standard normal distribution.

**Remark:** BONOBO derives a dense (complete) network with edges between every pair of genes, where edge weights correspond to *σ*_*jk*_, the posterior mean of the covariance between genes *j* and *k*. We can generate a sparse covariance network by simply pruning out edges for which the 100(1−*α*)% posterior credible regions contain zero, for a suitable value of 0 *< α <* .

## 3 Results

### 3.1 Simulated Data and comparison with other methods

Although we recognize that simulated gene regulatory networks may not capture the full complexity of the gene expression and the effects of regulation, simulation is an important tool as it provides a measure of “ground truth” against which various methods can be rigorously benchmarked and compared. We performed five simulation experiments and compared BONOBO with two other methods for computing sample-specific co-expression: LIONESS [9] and SPCC [7, 8]. We repeated each of the following simulation experiments 100 times and compared BONOBO with LIONESS and SPCC based on the mean sum of squared errors (i.e. the squared Frobenius distance between the true correlation matrix and the estimated correlation matrix) across these 100 iterations. Through samples of varying sizes and different dimensionalities from a homogeneous population we demonstrate that (i) increasing sample size improves the performance of BONOBO at a faster rate than other methods (Supplementary Materials S1.2.1) while (ii) increasing the number of genes deteriorates the performance of all three methods (Supplementary Materials S1.2.2). Next, (iii) we simulated samples from a mixture of two different homogeneous populations (Supplementary Materials S1.2.3) and demonstrated that the performance of BONOBO remains unaffected by the mixing proportion of the two populations. In all these instances of simulation experiments, the mean squared errors for BONOBO were much smaller than those for LIONESS and SPCC. In the next example, (iv) we simulated samples from a mixture of two populations, where one population lost expression for 1% of genes. BONOBO identified this loss of expression (Supplementary Materials S1.2.4) with better accuracy, compared to the two competing methods. Finally (v) we used two simulation examples to illustrate (Supplementary Materials S1.2.5) that the data-driven approach of calibrating hyperparameter *δ* described in section 2.1.1, provides comparable estimation accuracy, compared to the optimal performance obtained by choosing an arbitrary fixed value of *δ*.

Taken together, we have simulated datasets that that resemble many of the scenarios encountered in biological datasets such as different population sizes, single gene knockouts and silencing, mixtures of subpopulations. In all these conditions, BONOBO performs better than both LIONESS and SPCC, thereby demonstrating the efficacy of our method in capturing the “true” correlation patterns between genes, compared to the existing methods for sample-specific coexpression estimation.

### 3.2 BONOBO recovers sample-specific network structure in yeast datasets

Having established the performance characteristics of BONOBO for simulated data, we want to prove that our method is applicable and useful on experimental data. *Saccharomyces Cerevisiae*, i.e. yeast, is a well-studied organism and it has been extensively used to model biological networks [18, 19, 20]. Moreover, perturbation experiments have been key to study the connectivity between biological entities in yeast [21, 22, 23]. We reason that using yeast experiments would allow us to test BONOBO on real data, validating our findings with those in the literature, and testing our method’s ability to detect perturbations at the single sample level. First, we applied BONOBO to 48 cell-cycle-synchronized yeast microarray samples [24]. With this smaller dataset we assess the behavior of p-value thresholding in real data and we show that by using BONOBO networks we are able to detect the fluctuations in the cell-cycle transition pathways (see Supplementary Materials S1.3).

Thus, we applied BONOBO to a yeast gene perturbation experiment [25] that includes pseudo-bulked gene expression from 132 engineered strains that combine genetic and environmental perturbations; 11 TF knockout (KO) genotypes target the Nitrogen Catabolite Repression (NCR) pathway, the General Amino Acid Control (GAAC) pathway, the Ssy1-Ptr3-Ssy5-sensing (SPS) pathway, and the retrograde pathway. These strains were grown in twelve conditions that included various nitrogen and carbon sources (Supplementary Materials S1.4). While this is originally a scRNA-seq dataset, we created pseudo-bulk expression values for each KO-media combination. This way we can use BONOBO to generate networks that are perturbation-specific and investigate the co-expression changes induced by each KO and media.

Consistent with the results of the original paper, we find that BONOBO’s networks in the same growth medium tend to be more similar than those with the same KO (Figure 2A). This makes also logical sense, since different nutrients will perturb larger metabolic pathways, rather than one TF and its gene targets as it is the case for the TF KOs. However, leveraging BONOBO’s sample-specific co-expression, we can analyze how much the gene deletion affects each sample’s network by looking at the effects of a specific TF KO on the other genes and comparing it to the other samples (Figure S5). It appears that deletion of GCN4 has the strongest effect on the network, visibly changing the correlation patterns of the genes that have the highest edge values with GCN4 (Figure S6). Furthermore, we can pinpoint the effects of GCN4 deletion on each network by reconstructing which edges are the most affected by the perturbation. We then selected the top 100 genes whose edge with GCN4 is the most affected by the perturbation, by means of checking which edges are most different between the GCN4 genotype and the rest of the dataset. As expected, genes that are most perturbed by GCN4 deletion, that targets the GAAC pathway, belong to many pathways related to autophagy such as exosome, phagosome, and autophagy - yeast (Figure 2B). Interestingly, between the top perturbed genes there are known GCN4 targets [26] such as VCX1, MNN10, ACT1,CPA1 ([27, 28], and CCW12 [29] showing that by estimating perturbation-specific co-expression networks BONOBO is able to detect key interactors of the TF. The analysis of yeast experiments recapitulate many of the known properties of yeast cells, showing that BONOBO works well on real data, in addition to simulated datasets. Moreover, BONOBO allows us to investigate the effects of each combination of TF KO and growth media, which would be impossible with conventional population-based correlation measures.

**Figure 2:**
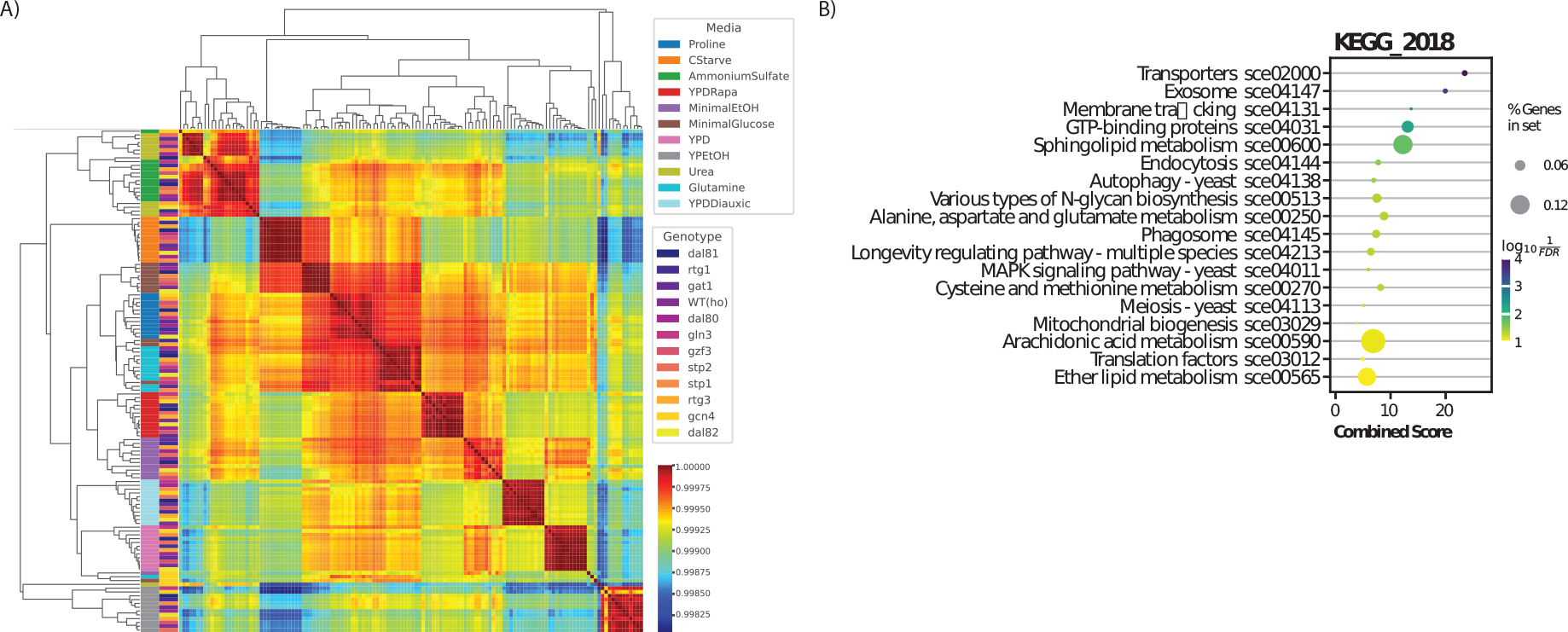
A) Pearson correlation between all sparsified BONOBOs computed on the engineered yeast data. Row labels report both the gene deletion and the growth medium for each sample. As expected, co-expression networks tend to correlate more with the strains subject to the same growth condition, rather than those with the same deletion.B) over-representation analysis of the top 50 genes disrupted by GCN4 perturbation. The enriched terms belong to the KEGG 2018 pathways.

### 3.3 miRNA-gene Interaction in Breast Cancer Subtypes

As evidenced from the simulation experiments and the analysis of gene expression data from perturbed yeast cells, BONOBO successfully recovers sample-specific heterogeneity in the gene co-expression networks. Furthermore, we propose that BONOBO can serve as a useful tool for investigating interaction patterns between multiple omics modalities, while accounting for sample-specific clinical and molecular confounders. MicroRNAs, or miRNAs have been observed to play important parts in RNA silencing by down-regulating the expression of multiple genes and modifications in miRNA levels have been shown to be involved in the development and prognosis of various cancer types [30]. Here, we used individual sample-specific co-expression networks constructed using paired mRNA and miRNA bulk expression data from multiple breast cancer subtypes (GEO accession number GSE19783 [31, 32, 33], preprocessing steps in Supplementary Materials S1.5.1) to understand how correlation between miRNA and genes (or biological pathways) vary across different breast cancer subtypes and whether this interaction between genes and miRNAs have any association with breast cancer survival.

In total we have 101 sample-specific co-expression networks each of which constitutes of correlations between pairs of genes, pairs of miRNAs and between each gene and each miRNA. From these BONOBO networks we are interested in investigating which biological pathways are most significantly associated with miRNA expression in various breast cancer subtypes. For each network, we account for the association between a gene and all miRNA by summing over all edges connecting the particular gene to the miRNAs and we then applied pathway analysis (Supplementary Materials S1.5.2).

We observed that pathways associated to immune response including Graft vs Host disease, primary immunodeficiency, cytokine-cytokine receptor interaction and natural killer cell mediated cytotoxicity were significantly (at FDR cutoff 0.05) negatively correlated with miRNA expression, across all breast cancer subtypes (Figure S7). Pathways associated to cell adhesion and cell proliferation, such as focal adhesion and ECM receptor interaction were positively correlated with miRNA expression, especially in Basal, Normal-like and Luminal B subtypes, while pathway associated to cell adhesion molecules was significantly (at FDR cutoff 0.05) negatively correlated with miRNA expression in ERBB2, Luminal A and Normal-like breast cancer. These findings align with previous studies [31] that also uncovered significant associations between the expression levels of several miRNAs and biological pathways involved in cell proliferation, cell adhesion and immune response.

Although pathways most correlated (positively or negatively) with miRNA expression were consistent across different breast cancer subtypes, upon closer inspection of the individual mRNA-miRNA edges in the networks, we observe that there is significant difference in the neighborhood of individual genes across different breast cancer subtypes. In particular, comparing individual edges between genes and miRNAs in samples from luminal A and luminal B (Figure 3B) subtypes, we found several genes that were differentially correlated with certain miRNAs in luminal A versus luminal B subtypes. Genes most differentially coexpressed with miRNAs include proto-oncogene *EGFR*; genes involved in immune response such as *PI3KCD and INFGl*; genes involved in cell-cell adhesion and cell proliferation such as *CLDN1, CLDN8, CLDN10, CLDN16* etc. These differences between miRNA-gene interactions highlight the distinct molecular landscapes of luminal A and luminal B, which might be a contributing factor towards diverse clinical presentations observed in both the incidence rates and prognoses of these two breast cancer subtypes. Previous research [34] has also demonstrated that the miRNA dysregulation patterns in luminal A breast cancers differ from those in luminal B breast cancers.

**Figure 3:**
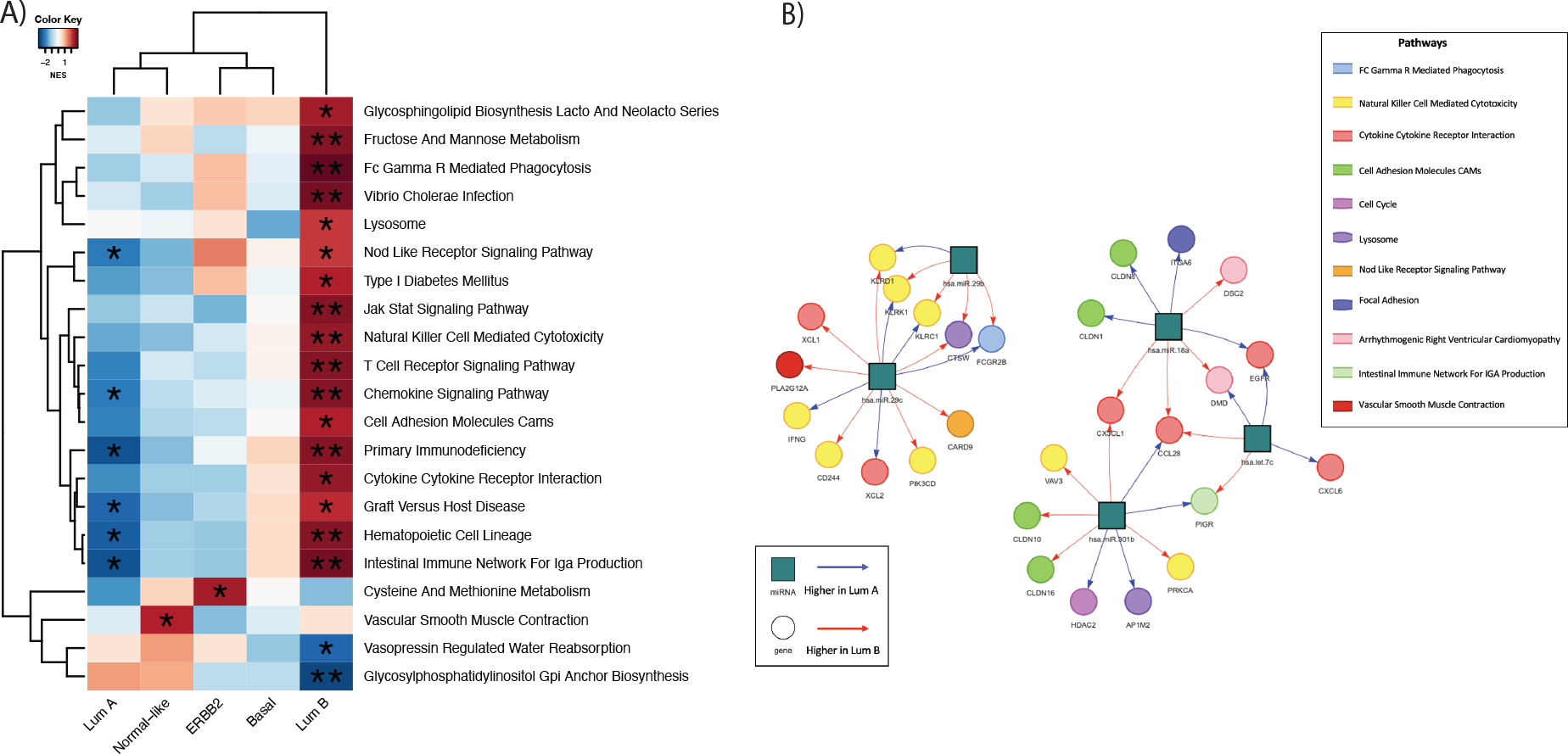
A) Pathways for which the mean correlation with miRNAs is significantly (at significance level 0.05) associated with survival in Cox proportional hazard model: heatmaps are colored by t-statistic of the pathway score in the Cox model; Pathways for which higher correlation with miRNAs is associated to better survival are colored blue and pathways for which higher correlation with miRNAs is associated to worse survival are colored red. B) Pairs of genes and miRNAs that are most differentially coexpressed between Lumina A and Luminal B subtypes: genes nodes are represented by colored circles where the colors correspond to the the biological pathway associated to the gene.

Next we investigated, if the strength of association between miRNA expression and biological pathways have any influence on the survival outcome in various breast cancer subtypes. For every pathway significantly (at FDR cutoff 0.05) correlated (positively or negatively) with miRNA expression, we computed a pathway score, defined by the mean of the total correlation between all miRNAs and each gene in that pathway. Then for every pathway, we fit a Cox proportional hazard model to predict survival, using the pathway scores, while allowing the coefficient of the Cox model to be subtype-specific by including an interaction effect between the pathway score and the breast cancer subtype of every sample.

We observe that a higher correlation between miRNA and pathways associated with immune response such as Chemokine signaling pathway, primary immunodeficiency, Graft vs Host disease, Hematopoietic cell lineage and intestinal immune network for IGA production was associated with better survival among samples from luminal A subtype, while having worse survival among samples from luminal B subtype (Figure 3A). Previous studies [35] have also demonstrated that an increased expression of miRNAs is associated with tumor suppression among luminal A breast cancers, thus leading to slower tumor growth and improved prognosis. Our analysis indicates that an up-regulation of immune pathways by miRNAs might provide a possible mechanisms for tumor suppressive effect of miRNAs in luminal A. In contrast, in luminal B breast cancer, miRNAs may have a role in promoting immune evasion [36], thus leading to more aggressive tumor growth and poorer survival outcome.

Previous studies [37] had identified specific immune gene expression patterns that distinguish luminal A from luminal B subtypes and showed that these distinct immune signatures were associated with a differential ratio between ESR1 and ESR2, a higher value of which was further associated to poorer survival outcome. Our results indicate that subtype-specific regulatory interactions between miRNAs and immune pathways in luminal A versus luminal B breast cancers might be a possible factor contributing towards estrogen receptor mediated survival outcome. Furthermore, we also demonstrated that the genes linked to cell proliferation pathways exhibit distinct patterns of regulation by miRNAs in luminal A versus luminal B breast cancer subtypes. Clinically, this disparity in the regulation of cell proliferation genes might be a crucial factor contributing to the worse prognosis associated with luminal B breast cancer, as it tends to exhibit heightened cellular proliferation [38]. In conclusion, individual-specific heterogeneity in correlation networks between miRNAs and genes can provide valuable insights into the distinct miRNA-gene interaction patterns that distinguish various breast cancer subtypes, which in turn might have significant implications for breast cancer prognosis and personalized therapeutic strategies.

### 3.4 Sample-specific Gene Regulatory Networks Identify Sex Difference in Thyroid

Most thyroid disorders have sex difference in incidence rate with females being significantly more susceptible to be affected by a thyroid condition at some point in their lives, compared to males [39]. Although sex chromosomes, sex hormones, and the immune system [40] have often been cited as possible contributors to this sex difference in thyroid tissue, a system-based analysis exploring the regulatory mechanisms associated to these sex-biased disease manifestation is scarce. We used sample-specific gene regulatory networks constructed from healthy thyroid tissue samples from the Genotype Tissue Expression (GTEx) Project [41] (see Supplementary materials S1.6.1 for all preprocesing steps), to understand how the genes are differentially regulated by transcription factors (TF) in males and females. We combined the sample-specific co-expression networks derived by BONOBO with TF-motif prior information (Supplementary materials S1.6.2) and protein-protein interaction data (Supplementary materials S1.6.3), using the PANDA network inference algorithm[3] to derive bipartite sample-specific gene regulatory networks that connect transcription factors (TFs) to their target genes (Figure S8).

Based on the differential targeting analysis (Supplementary materials S1.6.4), we observed that several genes with known relevance in various thyroid cancers and autoimmune conditions are differentially targeted by transcription factors in males and females (Figure S10). The long non-coding RNA *XIST* showed higher targeting in females. Previously, *XIST* had been observed to promote oncogenic activities in papillary thyroid carcinomas (PTC) [42]. Among genes highly targeted in males, tumor suppresor gene *KDM6A* is known to regulate multiple genes involved in immune response, suggesting its potential influence on the risks of developing various autoimmune conditions [43]; Among other genes highly targeted in males, *PCM1* mutation has also been associated with PTC [44]; while mutation in *KMT2C* has been identified as a molecular marker for primary thyroid osteosarcoma [45]. Additionally, overexpression of *SOS1*, which also showed higher targeting in males, have been found to promote cell proliferation and cell apoptosis in PTC cells [46].

Finally, we functionally characterised the genes differentially targeted in male and female samples (see Supplementary materials S1.6.5). We observed that biological pathways associated to immune response such as humoral immune response, B-cell receptor signaling pathway, antigen-receptor mediated signaling and positive regulation of b-cell activation pathway were targeted more in females (Figure 4). Heightened targeting of immune pathways in females may contribute to their increased susceptibility to autoimmune thyroid diseases including Hashimoto’s Thyroiditis disease, which often lead to hypothyroidism. On the other hand, pathways associated to cell cycle, cell signaling, metabolic processes and DNA repair were targeted more in males (Figure 4). Disregulation of these pathways have been shown to play integral parts in various thyroid conditions including Graves’ Disease and Hashimoto’s Thyroiditis [47] and therefore the differential targeting of these pathways in males suggests that they may have a more robust defense against factors that could disrupt thyroid function or lead to the development of these thyroid diseases.

**Figure 4:**
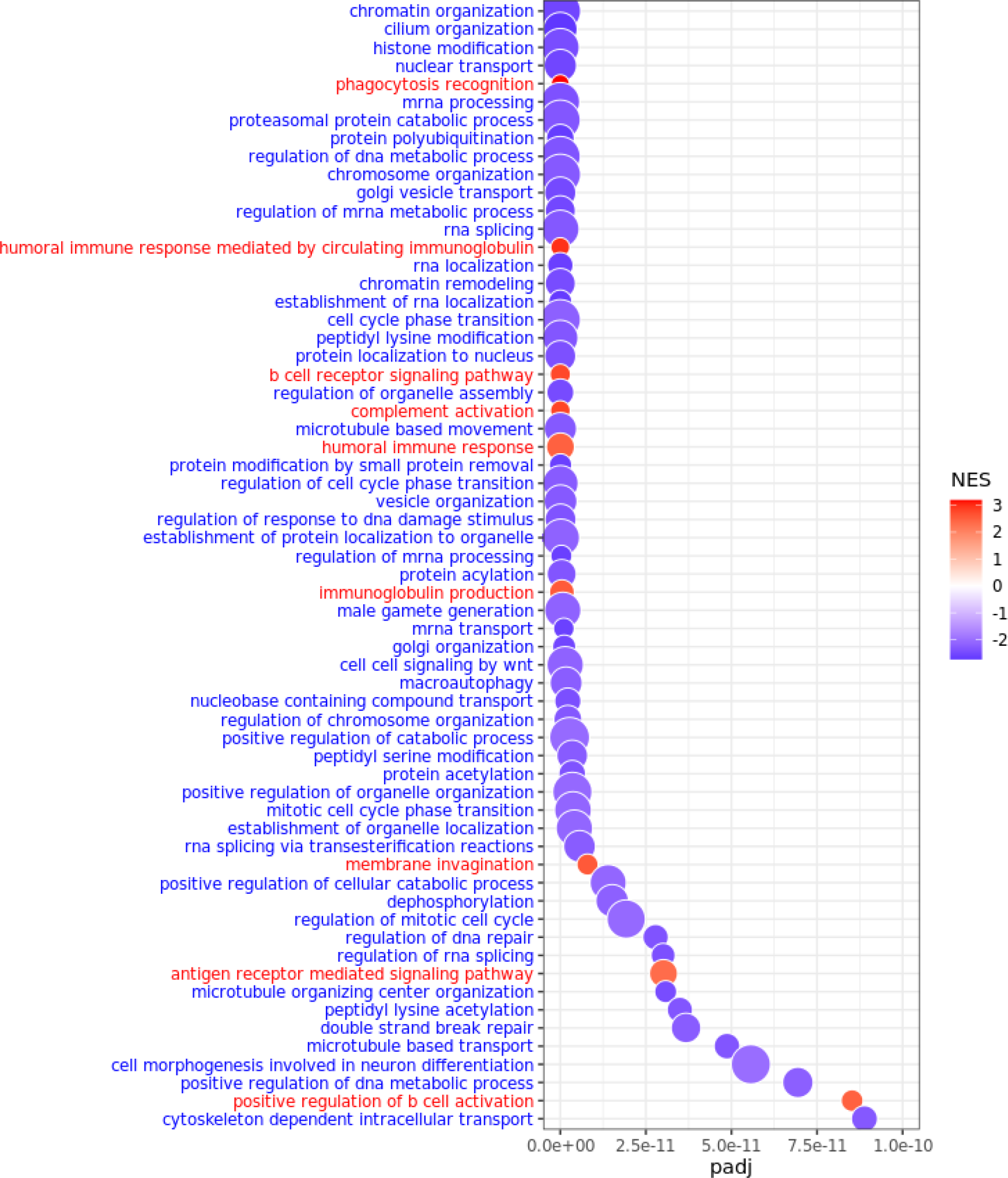
GO biological processes most differentially regulated (at FDR cutoff 1e-10) in males and females in GTEx thyroid samples: pathways highly targeted in males are marked in blue and pathways highly targeted in females are marked in red.

In conclusion, sex-biased differential regulation of key genes and biological pathways might be a contributing factor towards differential risk of various thyroid conditions across both sexes and deciphering these sex-specific gene regulatory patterns through individual-specific gene regulatory networks can aid in developing more effective, personalized interventions for the prevention and treatment of various thyroid diseases.

## 4 Conclusion

Complex human traits and diseases are most often driven by not a single gene but rather by intricate interactions involving multiple genes and regulators. However, the majority of network inference techniques estimate an aggregate network [4, 6] that reflects the average regulatory characteristics of the population, thereby overlooking the diversity within the population that might arise due to various biological (e.g. sex-difference) and/or environmental factors (e.g. carcinogen exposure) [48]. Recognizing the inherent hererogeneity in regulatory processes among individuals, we have introduced BONOBO, a Bayesian parametric model designed to construct personalized sample-specific gene co-expression networks for single samples. These networks enable us to capture population heterogeneity in gene-gene interaction, that are rarely reflected in aggregate co-expression networks constructed by conventional algorithms.

Constructing individual-specific co-expression networks is particularly challenging in bulk expression data as each individual do not have more than one sample. Bayesian statistics enables us to overcome this dearth of individual-level data by incorporating prior information derived from other individuals in the same dataset. We assume that individual-level covariance matrices come from an inverse Wishart prior, whose mean is equal to the sample covariance matrix computed from all other individuals in the given dataset. This assumption is based on the fact that typically samples in a bulk expression dataset come from a single tissue and from individuals having similar conditions (e.g. primary tumor samples). Integrating this prior information with the individual-level expression data, we estimate posterior distribution of the covarince matrix for each individual in the data. Interestingly, the mean of the posterior distribution turns out to be a weighted average of the deviation of the individual expression from the mean expression and the estimated population covariance from all the other individuals in the dataset. Thus for every individual, BONOBO compensates for the lack of individual-level information by borrowing stength from all other individuals in the data.

BONOBO is highly scalable as we use conjugate prior distribution over individual-specific covariance matrices, which enables us to derive a closed form expression of the posterior distribution, thus eliminating the need for running computationally expensive Markov Chain Monte Carlo. In addition, the posterior distribution of the covariance matrix for each individual is computed separately, without any influence of the posterior distribution of other individuals, thus making BONOBO highly parallelizable, further enhancing computational efficacy.BONOBO is based on minimal assumptions and only one tuning parameter that can be efficiently calibrated using a data-driven approach. BONOBO assumes that for every individual in the data, the log transformed expression values of genes follow multivariate Gaussian distribution with a covariance matrix unique to every sample. Through simulated examples where the samples come from a mixture of two different populations with different mean expression and patterns of co-expression, we demonstrate that BONOBO performs better than competing methods even when the underlying assumption of multivariate normality is violated, thus making the method adaptable to a wide range of applications.

In addition to providing point-estimates for each pair of gene-gene correlation, BONOBO provides posterior credible intervals for these individual-specific correlation estimates. These credible regions enable the user to derive sparse co-expression networks by simply pruning out edges between pairs of genes that are uncorrelated with high probability. Using a perturbed yeast cell dataset, we illustrate that these sparse networks not only reflect the sample-specific perturbations globally, but they allow us to investigate the specific neighborhoods that are perturbed by transcription factor KO. We demonstrate that BONOBO can be readily extended to capture interactions between multiple omic data categories to derive individual-specific gene regulatory networks. As an example, we analyze BONOBO networks that capture individual-specific correlation structures between genes and miRNAs in various breast cancer subtypes. Consistent with existing literature [31] we find biological pathways associated to immune response and cell proliferation to be significantly correlated with miRNA expression. Furthermore, through survival analysis, we demonstrate that interactions between miRNAs and immune pathways have varying degrees of prognostic significance between lumina A and luminal B breast cancer subtypes.BONOBO can be combined with existing methods for estimating gene regulatory network, such as PANDA to derive individual-specific bipartite networks with directed edges from transcription factors to target genes. To demonstrate this, we use RNA-seq data from thyroid tissue samples in GTEx and compared the resulting networks between males and females to understand why thyroid conditions are more prevalent among females than males [40]. Our analysis reveals that biological pathways assocaied to cell proliferation, immune response, and various matabolic processes are differentially regulated between males and females, thereby providing a possible mechanism that might contribute towards the observed sex-disparity in the incidence rate of various thyroid conditions.

We also recognize that BONOBO has limitations, some of which could be addressed by future research. Although in this paper we applied BONOBO exclusively on transcriptomics data, the model can be readily adapted to uncover interactions between other omic data types (e.g. proteomics) with little to no modifications, as long as the data can be suitably transformed to resemble a unimodal distribution over a continuous support. However, it is important to note that BONOBO is not applicable for omics modalities characterized by binary or categorical data, such as mutation profiles. A promising avenue for future research could involve extending our model to incorporate interactions across a broader range of omics data types, through hierarchical latent variable models and/or association measures other than Pearson’s correlation. Extending BONOBO to other correlation measures, such as the Spearman’s rank correlation coefficient, would also allow to overcome Pearson’s correlation intrinsic limitations. For instance, Pearson’s correlation is heavily influenced by outliers, thereby potentially impacting all individual co-expression networks inferred by BONOBO. Moreover, Pearson’s correlation quantifies only the extent of linear association between genes and their molecular regulators. In future research we would like to extend BONOBO to estimate correlation networks based on Chatteejee’s correlation coefficient [49] since it is capable of capturing not only linear association but also functional dependence between pairs of genes.

In summary, we derive BONOBO, a Bayesian statistical model for deriving individual-specific co-expression networks, that can be further amalgamated with other network inference methods to infer individual-specific gene regulatory networks. BONOBO can be employed to capture population heterogenerity in interaction patterns involving multiple omics data types, thereby providing a more nuanced understanding of the complex mechanisms of human traits and diseases. Through various real datasets containing multiple omic modalities, we demonstrate that BONOBO can potentially enable network-based disease subtyping and facilitate individualized therapy design in diverse human diseases.

## Code Availability

BONOBO is available through the Network Zoo package (netZooPy v0.9.18; netzoo.github.io).

## Acknowledgements

This work was supported by grants from the National Institutes of Health: ES, CMLR, MBG, VF, JF, KHS, PM, and JQ were supported by R35CA220523; MBG and JQ were also supported by U24CA231846; JQ received additional support from P50CA127003; JQ and DLD were supported by R01HG011393; KHS and DLD were supported by R01HG125975 and P01HL114501; KHS was supported by T32HL007427; CMLR was supported by K01HL166376; CMLR and ES were also supported by the American Lung Association grant LCD-821824.

## Author Contributions

**Conceptualization:** ES, VF, PM, MBG, JF, KHS, KG, DLD, CLR and JQ; **Methodology:** ES and VF; **Formal Analysis:** ES, VF and PM; **Investigation:** ES and VF; **Resources:** JQ and MBG; **Data Curation:** ES, VF and CLR; **Writing – Original Draft:** ES and VF; **Writing – Review and Editing:** PM, MBG, JF, KHS, KG, DLD, CLR and JQ; **Visualization:** ES, VF and PM; **Supervision:** JQ, CLR, and DLD; **Funding Acquisition:** JQ, CLR, and DLD

