## Supplementary material for "Bayesian Optimized sample-specific Networks Obtained By Omics data (BONOBO)"

November 3, 2023

**S2 Supplementary Tables** **15**

### S1 Supplementary Materials and Methods

#### S1.1 Proofs of Theoretical results

##### S1.1.1 Proof of Theorem 1

Under assumption (1), the conditional likelihood of the centered expression of the $i$-th sample can be written as

$$\mathbb{P}(x_i - \bar{x}|V_i, \{e_j\}_{j\neq i}) \propto |V_i|^{-1/2} \exp\left(\frac{1}{2}(x_i - \bar{x})^T V_i^{-1}(x_i - \bar{x})\right) \tag{1}$$

Under assumption (2), given all other samples in the data, the prior probability density of the covariance matrix of the $i$-th sample, can be written as

$$\mathbb{P}(V_i|\{x_j\}_{j\neq i}) \propto |V_i|^{-(\nu_i+g+1)/2} \exp\left(\frac{1}{2} trace\left((\nu_i - g - 1)S_i V_i^{-1}\right)\right) \tag{2}$$

Combining 1 and 2, the posterior distribution of $V_i$ can be written as

$$\begin{aligned}
\mathbb{P}(V_i|\{x_i\} \cup \{x_j\}_{j\neq i}) &\propto \mathbb{P}(x_i - \bar{x}|V_i, \{x_j\}_{j\neq i})\mathbb{P}(V_i|\{x_j\}_{j\neq i}) \\
&\propto |V_i|^{-(\nu_i+g+1+1)/2} \exp\left(\frac{1}{2}\left[(x_i - \bar{x})^T V_i^{-1}(x_i - \bar{x}) + trace\,(\nu_i - g - 1)\, S_i V_i^{-1}\right]\right) \\
&\propto |V_i|^{-(\nu_i+g+1+1)/2} \exp\left(\frac{1}{2} trace\left(\left[(x_i - \bar{x})^{-1}(x_i - \bar{x})^T + (\nu_i - g - 1)S_i\right] V_i^{-1}\right)\right)
\end{aligned}$$

From the above expression we see that $V_i \sim InvWishart\left((\nu_i - g)\Sigma_i, \nu_i + 1\right)$, proving Theorem 1.

#### S1.2 Simulation Experiments

We compare BONOBO with LIONESS and SPCC through five simulation experiments. For each of the methods, the estimated sample-specific co-expression networks are compared with the true co-expression. We compute the mean squared error (MSE) across all gene pairs. We repeat each simulation experiment for 100 iterations (i.e., 100 different simulated datasets). Because LIONESS and SPCC do not bound the network edgeweights to values between $-1$ and $1$, the resulting MSE values are much higher than in BONOBO. Cosequently, we scaled the off-diagonal elements of each estimated co-expression matrix by the maximum of the absolute values of all the off-diagonal elements and we set the diagonal entries to one to resemble the Pearson correlation matrices produced by BONOBO, thus providing a more "fair" comparison.

##### S1.2.1 Simulation Example 1: Varying Sample Size

We selected the top 50 most variable and top 50 least variable genes from the GTEx thyroid samples. For this set of 100 genes, we computed the mean gene expression (denoted by $\mu_{100}$) and the covariance matrix of gene expression (denoted by $\Sigma_{100}$). Then we simulated the gene expression distribution of these 100 genes for $N$ (ranging from 10 to 200) individuals (samples) as follows: for every individual (sample) we simulated 100 independent observations from a 100-dimensional multivariate normal distribution with mean $\mu_{100}$ and covariance matrix $\Sigma_{100}$. The correlation matrix computed from these 100 observations were treated as ground truth for the gene co-expression corresponding to this individual. The mean of these 100 observations was then treated as a representative sample for the gene expression of that individual, to be used as input to BONOBO and other competing methods. We implemented BONOBO, LIONESS and SPCC on the resulting N gene expression samples to estimate sample-specific gene co-expression networks.

We observed (Figure S1) that as sample size increased from $N = 10$ to $N = 200$, the average MSE computed over 100 iterations of the simulation experiment, decreased rapidly for BONOBO and stabilized for sample size greater than $50$. For LIONESS and SPCC the average MSE did not vary much over different sample sizes. Moreover, irrespective of the sample size chosen, the mean MSE for BONOBO was much smaller than both the competitors.

##### S1.2.2 Simulated Example 2: Varying Network Size

We generate $N = 100$ individual gene expression samples for $G(= 100, 500, 1000)$ genes as follows. We selected the top $G/2$ most variable and top $G/2$ least variable genes from the GTEx thyroid samples. For this set of G genes, we computed the mean gene expression (denoted by $\mu_G$) and the covariance matrix of gene expression (denoted by $\Sigma_G$). Then we simulated the gene expression distribution of these G genes for 100 individuals (samples) by following the same procedure as described in Simulation Example 1 (section S1.2.1). From the average MSE (and standard deviation of MSE over 100 iterations) computed over 100 iterations (given in Table S2.1) for the three methods, we observed that irrespective of the number of genes, average MSE for BONOBO estimates were much smaller than those for LIONESS and SPCC.

Interestingly, we also observed that the mean MSE for both LIONESS and SPCC decreased as number of genes increased. However, upon closer inspection we saw that this decrease in MSE was driven by the fact that as more genes were included, the new gene pairs had lower correlation values on average than the correlations in the smaller gene sets, as evident from the mean correlation values for number of genes $G = 100, 500$ and $1000$ recorded in the last column of Table S2.1. So, we divide the MSE with the variance of correlation for each of the three setups ($G = 100, 500$ and $1000$) and record the scaled MSE in parantheses in Table S2.1. The scaled MSE increased for all three methods as the number of genes increased. The scaled MSE for BONOBO was much smaller than those for both LIONESS and SPCC, depicting better estimates of sample-specific co-expression matrices by BONOBO compared to the other methods.

##### S1.2.3 Simulated Example 3: mixture population

In this example we constructed a sample of 100 individuals $p\%$ ($p$ ranging from 0 to 50) of which come from one population and the remaining $(100 - p)\%$ come from a different population constructed as follows. We considered the same 100 genes as in Simulation Example 1 (section S1.2.1) from the GTEx thyroid samples. Then the GTEx thyroid samples were randomly partitioned into two groups so that one group contained $p\%$ of the samples and the other group contained the remaining $(100 - p)\%$. For each of the two partitions, we computed the mean gene expression of the 100 genes, denoted by $\mu_1$ and $\mu_2$. We generate 100 such random partitions and chose the partition with the highest Euclidean distance between the two means $(\mu_1 - \mu_2)^2$. The mean gene expressions $\mu_1$ and $\mu_2$ and the covariances of gene expression, denoted by $\Sigma_1$ and $\Sigma_2$ for these two partitions were then used as the parameters for simulation.

For each of the $p\%$ of samples, we simulated 100 observations from multivariate normal with mean $\mu_1$ and covariance matrix $\Sigma_1$. For each of the remaining $(100 - p)\%$ of samples, we simulated 100 observations from multivariate normal with mean $\mu_2$ and covariance matrix $\Sigma_2$. Finally, for every individual, the correlation matrix computed from the simulated 100 observations were treated as the ground truth for the gene co-expression corresponding to that individual. The mean of these 100 observations was then treated as a representative sample for the gene expression of that individual, to be used as input to BONOBO, LIONESS and SPCC.

From Figure S1 (middle), we see that as we increased $p$ from $0$ (all samples generated from the same population) to $50$ (equal number of samples generated from the two populations), irrespective of the value of $p$, the mean MSE for BONOBO was much smaller than both LIONESS and SPCC.

##### S1.2.4 Simulation Example 4: gene deletion

We considered the same 100 genes as in Simulation Example 1 (section S1.2.1) from the GTEx thyroid samples and generated the gene expression distribution for 100 individuals as described in Simulation Example 1. Then we randomly selected $p\%$ of individuals and $1\%$ of genes and set the gene expression value of the selected genes to 0 for these $p\%$ individuals, while keeping the other $(100 - p)\%$ individuals unaltered. For these $p\%$ of individuals all correlation values involving the $1\%$ selected gene would be exactly 0. Based on the mean MSE for the correlation values corresponding to these $1\%$ genes for the $p\%$ individuals (Figure S1 (right)), we see that for $p$ ranging between $5$ to $90$, average MSE for BONOBO computed over 100 iterations was lower than both LIONESS and SPCC. Although for $p$ greater than $90$, the mean MSE for all three methods were comparable.

##### S1.2.5 Simulation Example 5: varying degrees of freedom

We demonstrate on two simulated examples that computing prior degrees of freedom $\mu_i$, separately for every sample $i$, using the approach described in section 2.1.1, performs well, compared to assigning arbitrary fixed values of $\nu_i$ for all samples. Since $\nu_i$ is a one-one function of hyperparameter $\delta_i$, we equivalently compared the performance of BONOBO over varying values of $\delta$, where $\delta = \delta_i \forall i$.

In the first simulation example (Figure S2 (left)), we generate 100 random datasets, each containing 100 samples of gene expression values of 100 genes, as described in section S1.2.1.

In the second example (Figure S2 (right)), we generate 100 random datasets from a mixture distribution, where $20\%$ of the samples come from one population and the other $80\%$ samples come from another population with different mean expression. The mean of the larger population remained same as in S1.2.1, while the mean of the smaller population was $10\%$ higher for every gene, compared to the larger population. The covariance matrices for both populations were assumed to be the same, as in section S1.2.1.

For both examples, we computed the mean sum of squared errors (MSE) over these 100 datasets for each chosen value of $\delta_i = \delta$, ranging between zero and one. We then compared these MSEs with the MSE obtained from BONOBO using the data-derived calibrated values of $\delta_i$. We observed that in both simulation examples, the MSE for BONOBO with calibrated values of $\delta_i$ was comparable to the minimum MSE obtained by using a fixed choice of $\delta_i$ for all samples.

#### S1.3 Yeast Cell Cycle Data

We applied BONOBO to cell-cycle-synchronized yeast microarray data (Gene Expression Omnibus: GSE4987,[13]) that was previously normalized and pre-processed [12]. This experiment has microarray profiling of yeast cells, 2 cell cycles, made of 24 samples each (48 in total). The expression data shows periodicity of transcript levels, hence we expect the network to reflect these changes in cell-cycle phase. Indeed, by looking at the correlation between each couple of BONOBO networks, we find higher values in correspondence of the same cell cycle phase, across both replicates (Figure S3). Moreover, we applied the sparsification procedure for each BONOBO, with different confidence thresholds ($pval <$ $0.001$,$0.01$,$0.05$,$0.1$,$0.2$); while a low threshold such as $p < 0.001$ might be too restrictive and result in a low number of edges, we can see that the sparsification is reliable for values above 0.01. Interestingly, the number of non-zero edges for each of the samples captures the periodicity of the expression signal (Figure S3). At last, we apply a comparison method between BONOBOS in different cell-cycle phases. That is, we average the sparsified BONOBOS into three networks (G1, S, G2/M). By estimating the differences in degree between each node, we can run a Gene Set Enrichment Analysis (GSEA) on the ranked genes. Using the Gene Ontology database(downloaded from the Molecular Signatures Database (MSigDB) (http://www.broadinstitute.org/gsea/msigdb/collections.jsp)), we can see that both transitions are enriched for DNA replication processes. Reassuringly, while the G1 to S transition is enriched for positive G1/S regulation the S/G2-M transition is enriched for negative regulation of G1/S transition (Figure S4).

#### S1.4 Yeast Perturbation Dataset

We downloaded scRNA-seq data from [9] which assayed genetically and environmentally perturbed *Saccharomyces Cerevisiae* strains. The knockouts include TFs in the NCR, The Nitrogen Catabolite Repression (NCR) pathway (GAT1,GLN3,DAL80, DAL81, DAL82,and,GZF3), from the General Amino Acid Control (GAAC) (GCN4), from he Ssy1-Ptr3-Ssy5-sensing (SPS) pathway (STP1, STP2), and finally TFs from the retrograde pathway (RTG1, RTG3). Each of these strains were then grown in 11 media, with different carbon and nitrogen sources: Yeast Extract, Peptone, Glucose (YPD), YPD, Harvested after Post-Diauxic Shift (YPDDiauxic),YPD + 200 ng/mL Rapamycin (YPDRapa), Yeast Extract, Peptone, Ethanol (YPEtOH), Minimal Media, Glucose (MinimalGlucose), Minimal Media, Ethanol, (MinimalEtOH), Nitrogen Limited Minimal Media with Glutamine (Glutamine), Nitrogen Limited Minimal Media with Proline (Proline), Nitrogen Limited Minimal Media with NH4 (AmmoniumSulfate), Nitrogen Limited Minimal Media with Urea (Urea), Carbon Starvation (CStarve). For all strains we have scRNA-seq expression data, hence we created pseudo-bulk expression values by averaging the counts for all cells in each genotype-growth medium combination. Also, to avoid correlation inflation, we removed genes that have no counts in more than $20\%$ of samples. In total, we generate BONOBOs for $132$ samples, and we compute the co-expression networks on $8804$ genes. Each sample name reports the gene perturbation and medium condition, for instance, sample 'gcn4_ypd' refers to the sample where gene GCN4 is knocked-down and the strain is grown in YPD. Finally, pathway overrepresentation analysis is conducted with the GSEApy package [6] using the KEGG 2018 gene set.

#### S1.5 miRNA-mRNA co-expression in breast cancer subtypes

##### S1.5.1 Data preprocessing

Expression profiling of $489$ miRNAs along with genome-wide matched mRNA profiling from $41046$ probes in 101 human primary breast tumor samples were obtained from the Gene Expression Omnibus (GEO) repository (GEO accession number GSE19783 [5, 1, 8]). Expression data along with clinical information were downloaded from GEO using R package GEOquery (version 2.62.2). The samples belonged to either five breast cancer subtypes. Three samples with unspecified subtype, along with three samples with missing subtype information were categorized as "not classified" for downstream analysis. Thus, for downstream analysis we consider 6 categories of subtypes: (i) Basal like or triple negative (n = 15); (ii) ERBB2 or Her2 positive (n=17); (iii) luminal A (n=41); (iv) luminal B (n=12); (v) Normal-like (n=10) and (vi) not classified (n=6).

From the expression data we removed mRNA and miRNA probes with missing values for all samples, thus leaving 41002 mRNA probes and 495 miRNA probes. We further removed unannotated miRNAs. For genes with multiple probes, we kept only the probe with the maximum variance and discarded the rest. From the $19597$ unique genes, we further removed $74$ Y genes and $19$ unmapped genes. We constructed sample-specific correlation networks with BONOBO using the remaining $19504$ genes (including $738$ X genes) and $384$ miRNAs. This resulted in 101 sample-specific co-expression networks each of which constitutes of correlations between pairs of genes, pairs of miRNAs and between each gene and each miRNA.

##### S1.5.2 Differential Coexpression Analysis

To identify which biological pathways are most significantly associated with miRNA expression in various breast cancer subtypes, we first computed a miRNA-specific in-degree of all genes, separately for each sample-specific BONOBO network, by summing over all edges connecting the particular gene to the miRNAs. Then we fit linear regression on the in-degree of all genes using the breast cancer subtype-specific intercepts, using R package "limma" (version 3.50.3) [14]. For every cancer subtype, genes were ranked by the t-statistic of the "limma" model and a gene set enrichment analysis was performed using gene sets from the Kyoto Encyclopedia of Genes and Genomes (KEGG) pathway database [10] ("c2.cp.kegg.v2022.1.Hs.symbols.gmt"). Multiple testing corrections were performed using the Benjamini-Hochberg procedure [3].

#### S1.6 Sex-differences in thyroid gene expression

##### S1.6.1 Preparing Gene Expression Data for Analysis

RNA-Seq data of 706 healthy thyroid tissue samples from the GTEx Project were downloaded from the Recount3 database [19] using R package "recount3" (version 1.4.0). Clinical data for GTEx samples were accessed from the dbGap website (https://dbgap.ncbi.nlm.nih.gov/) under accession number phs000424.v8.p2. We removed 53 samples because they were designated as "biological outliers" in the GTEx portal for various reasons (as described in https://gtexportal.org/home/faq). The remaining 653 samples (434 males and 219 females) were used in the final analysis.

Gene expression data were normalized by Transcripts per Million (TPM), using the "getTPM" function in the Bioconductor package "recount" (version 1.20.0) [4] in R (version 4.1.2). Lowly expressed genes (with counts $\leq 1$ TPM in at least $10\%$ of the samples) were filtered out, thus leaving 26792 (including 64 Y genes) genes for analysis. To build gene regulatory networks, we kept only those genes (26507 genes) that were present both in the filtered gene set, as well as in the TF/target motif priors (see sections S1.6.2 and S1.6.3).

We verified that the self-reported gender for the GTEx samples aligned perfectly with the biological sex through a principal component analysis of gene expression values of 64 genes on the Y chromosome [Figure S9]. Since some genes on the Y chromosome were assigned expression values in females due to mismapping of transcripts, we manually set Y chromosome gene expression values to zero for biological females.

Finally log2-transformed TPM normalized gene expression data were used as input to BONOBO to derive sample-specific co-expression networks. These co-expression networks along with sex-specific TF/target gene regulatory prior (obtained by mapping TF motifs from the Catalog of Inferred Sequence Binding Preferences (CIS-BP) [18] to the promoter of their putative target genes) and protein-protein interaction prior (using the interaction scores from StringDb v11.5 [16] between all TFs in the regulatory prior) were used as input in the PANDA algorithm to derive sample-specific gene regulatory networks, using python package netZooPy (version 0.9.10) [2].

###### S1.6.2 Designing Sex-specific Transcription Factor-Gene Motif Prior

The TF-gene motif prior network is a binary network that features edges connecting transcription factors (TFs) to their target genes, with the edge values (0 or 1) indicating the presence or absence of a transcription factor motif within the promoter region of a specific target gene. To create this regulatory motif network, we first obtained transcription factor motifs for Homo sapiens with direct or inferred evidence from the Catalog of Inferred Sequence Binding Preferences (CIS-BP) Build 2.0, accessible at http://cisbp.ccbr.utoronto.ca. These transcription factor position weight matrices (PWM) were then mapped to the human genome (hg38) using FIMO [7]. We retained only the highly significant matches ($p \leq 10^{-}5$) occurring within the promoter regions of Ensembl genes (specifically, Gencode v39 annotations retrieved from http://genome.ucsc.edu/cgi-bin/hgTables). These promoter regions were defined as the interval of [-750; +250] base pairs centered around the transcription start site (TSS). This comprehensive process yielded an initial set of potential regulatory interactions, involving 997 transcription factors that collectively targeted 61,485 genes.

To enable statistical comparisons between networks, we needed to ensure that male and female motif networks had identical sets of edges. In this regard, we built sex-specific transcription factor regulatory priors to account for the absence of Y chromosome genes in females. Within the female-specific regulatory prior, edges originating from or connecting to Y chromosome genes were effectively assigned a weight of zero, thus resulting in a network consisting of 52,266 edges. These same sex-specific TF-gene regulatory priors were also used in an earlier study to identify sex difference in regulatory processes associated with lung adenocarcinoma [15].

###### S1.6.3 Designing Protein-protein Interaction Prior

We obtained PPI data from the StringDB database (version 11.5) using the STRINGdb Bioconductor package [17]. Subsequently, we filtered the PPI data to retain only interactions between transcription factors in the TF-motif network (using a score threshold index of 0). To maintain consistency in PPI scores, we normalized them by dividing each score by 1000, thereby restricting the values to a uniform range of 0 to 1 for both the PPI dataset and the TF-motif network. Additionally, we set self-interactions between transcription factors to a value of one. Since PPI networks are inherently undirected, we transformed the data into a symmetric PPI matrix.

###### S1.6.4 Differential Targeting Analysis using Sample-specific Regulatory Networks

For every sample-specific gene regulatory network, we calculated the targeting score for each gene, equivalent to the gene's in-degree (defined as the sum of all incoming edge weights originating from all TFs within the network). Gene targeting scores between males and females were compared using linear regression models using R package “limma” (version 3.50.3). The linear models accounted for the effects of relevant confounders, such as sex (Male and Female), race (White, Black or African American, Others and Unknown), age, smoking status (ever-smoker and never-smoker), ischemic time, RNA integrity number and batch.

##### S1.6.5 Pathway Analysis of differentially targeted genes in thyroid samples

We performed Gene Set Enrichment Analysis (GSEA) using R package “fgsea” (version 1.20.0) [11] and gene sets from the Gene Ontology Biological Processes (GOBP) downloaded from the Molecular Signatures Database (MSigDB) (http://www.broadinstitute.org/gsea/msigdb/collections.jsp). Only gene sets of sizes greater than 15 and less than 500 were considered, after filtering out genes that are not present in the expression dataset. Genes were ranked by the t-statistics produced by the limma differential targeting analysis. Multiple testing corrections were performed using the Benjamini-Hochberg procedure [3].

### Supplementary Figures

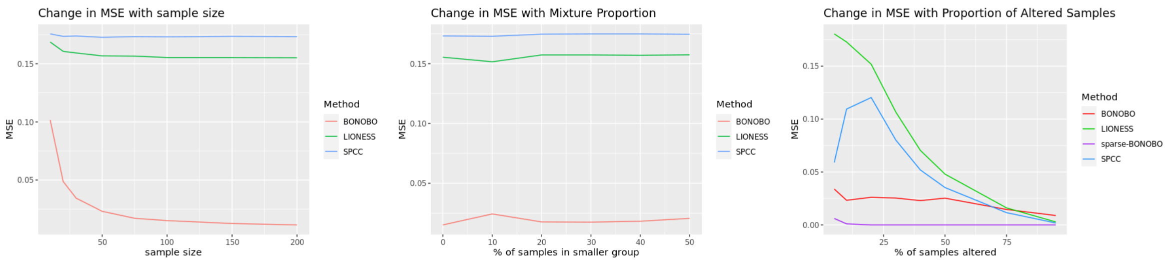

Figure S1: Performance of BONOBO, LIONESS and SPCC on simulated data: (Left) Simulated data from homogeneous population: Change in MSE with respect to sample size; (Middle) Simulated data from mixture of two populations: Change in MSE with respect to percentage of samples in the smaller population; (Right) Simulated data where some samples lost expression of 1% genes : Change in MSE with respect to the proportion of altered samples.

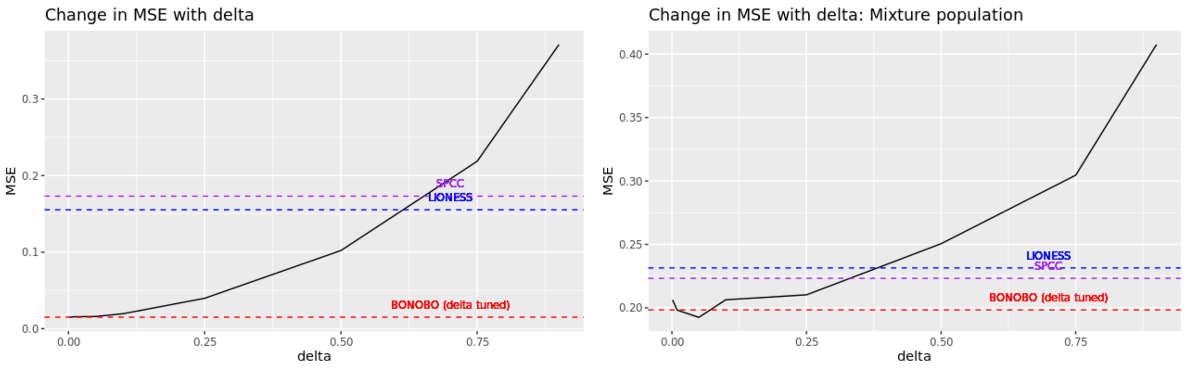

Figure S2: Change in MSE of BONOBO for varying values of hyperparameter $\delta$. MSE for BONOBO networks computed using $\delta$ estimated from the data are shown in red. MSE for LIONESS and SPCC are shown in blue and purple respectively. (Left) Simulated data from homogeneous population; (Right) Simulated data from mixture of two populations.

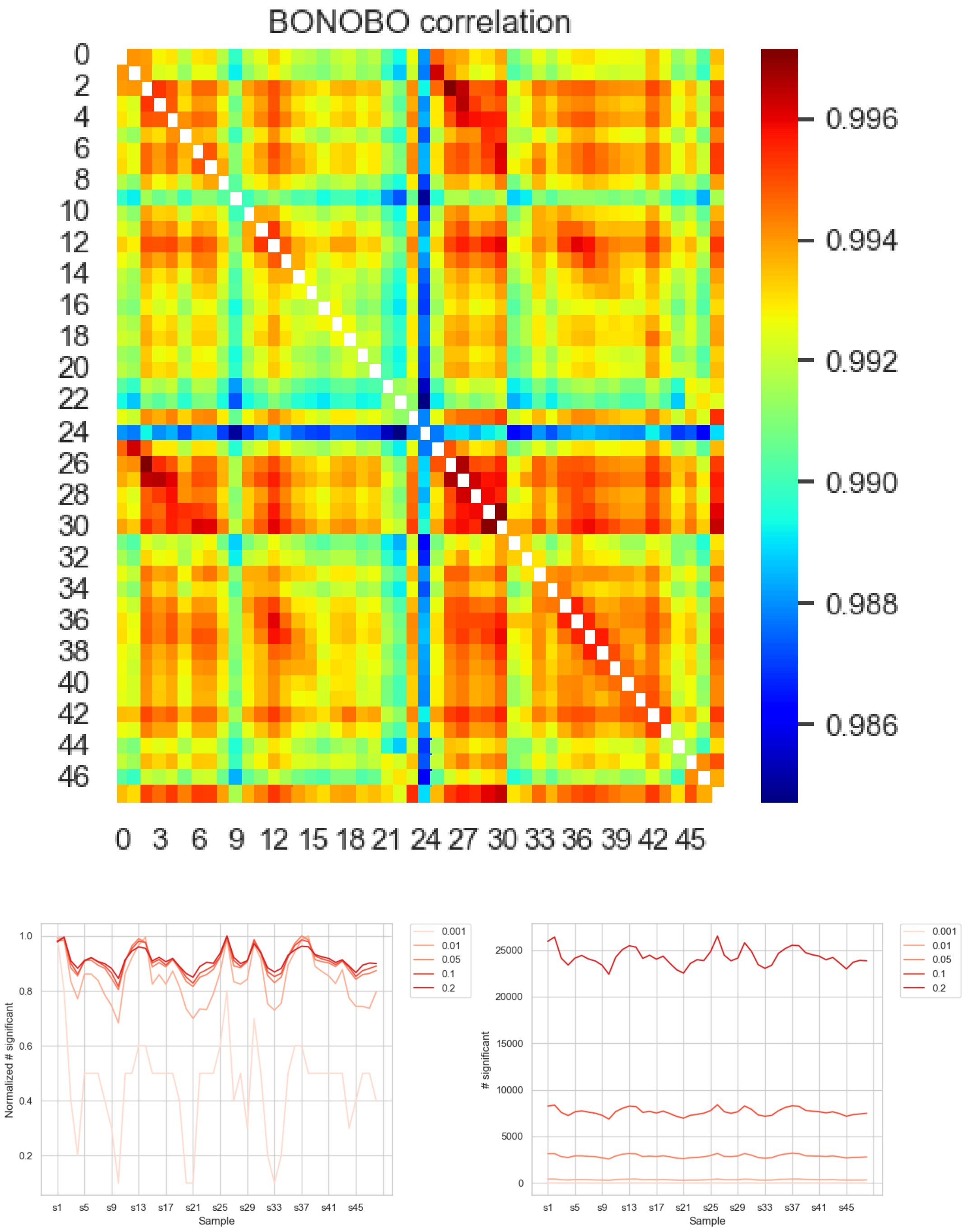

Figure S3: Top: Correlation between BONOBO network for all 48 yeast cell-cycle samples. Bottom: Proportion (right) and number (left) of non-zero edges for all sparsified BONOBO

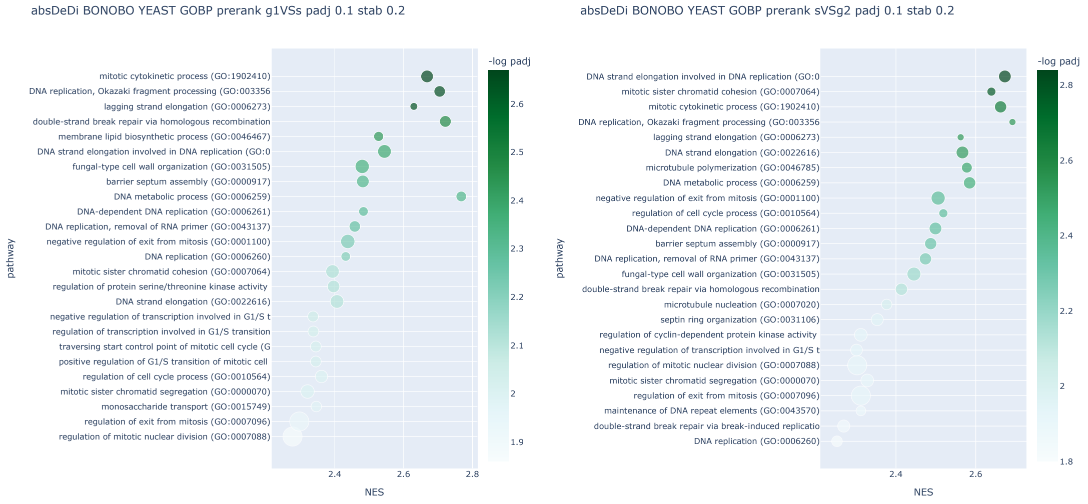

Figure S4: GSEA analysis of the G1 to S network comparison (left) and the S to G2/M networks (right). For each gene we compute the degree and then we compute the absolute degree difference (absDEDI) between two averaged networks. The genes ranked by absDEDI are used for the GSEA, and tested for all gene ontology biological processes.

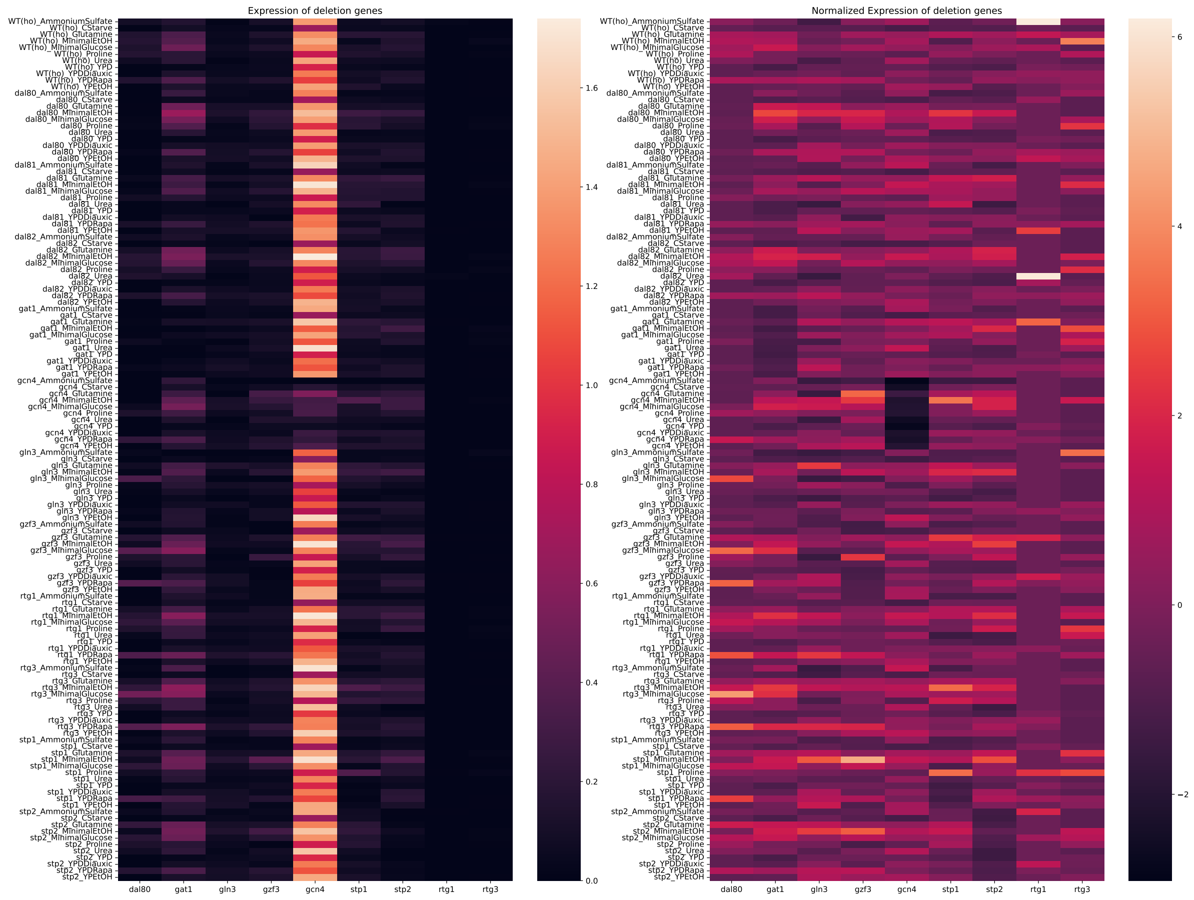

Figure S5: Expression of the perturbed TFs in each of the strains. On the left we have raw pseudobulk counts, on the right we have normalized the values by columns, to make them visually comparable. We can see that GCN4 is on average the most expressed TF, while also being the one for which the effect of deletion is better visible

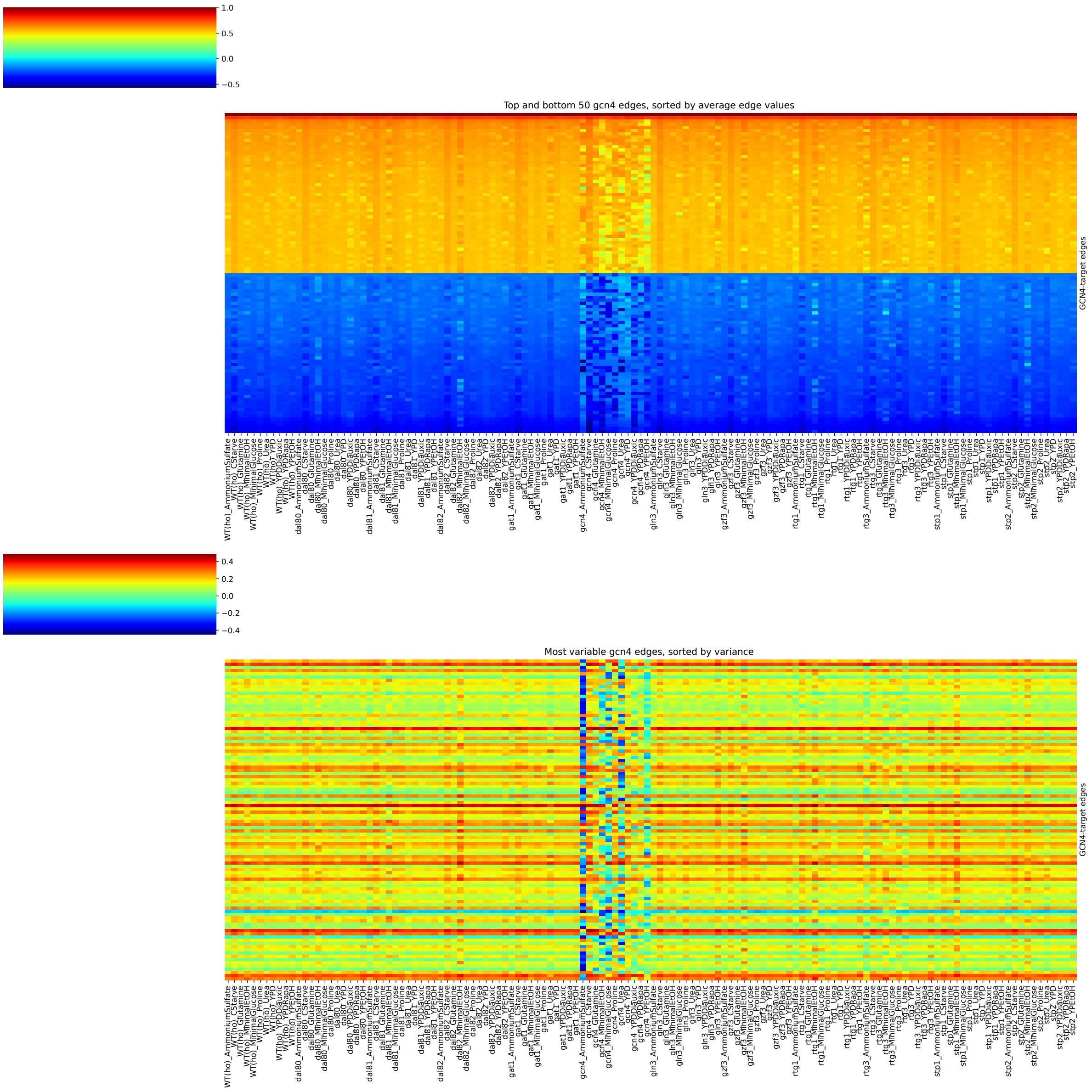

Figure S6: Top: strongest edges between GCN4 and any other gene. Genes are sorted by average mean value of the edges (row-wise), and then we selected the top and bottom 50 genes. Bottom: most variable edges between GCN4 and any other gene. Genes are sorted by average variance value of the edges (row-wise), and then we selected the top 100 genes by variance. In both cases we can see that the effect of GCN4 knockout (central columns of the heatmaps) produced an effect on the edge weights.

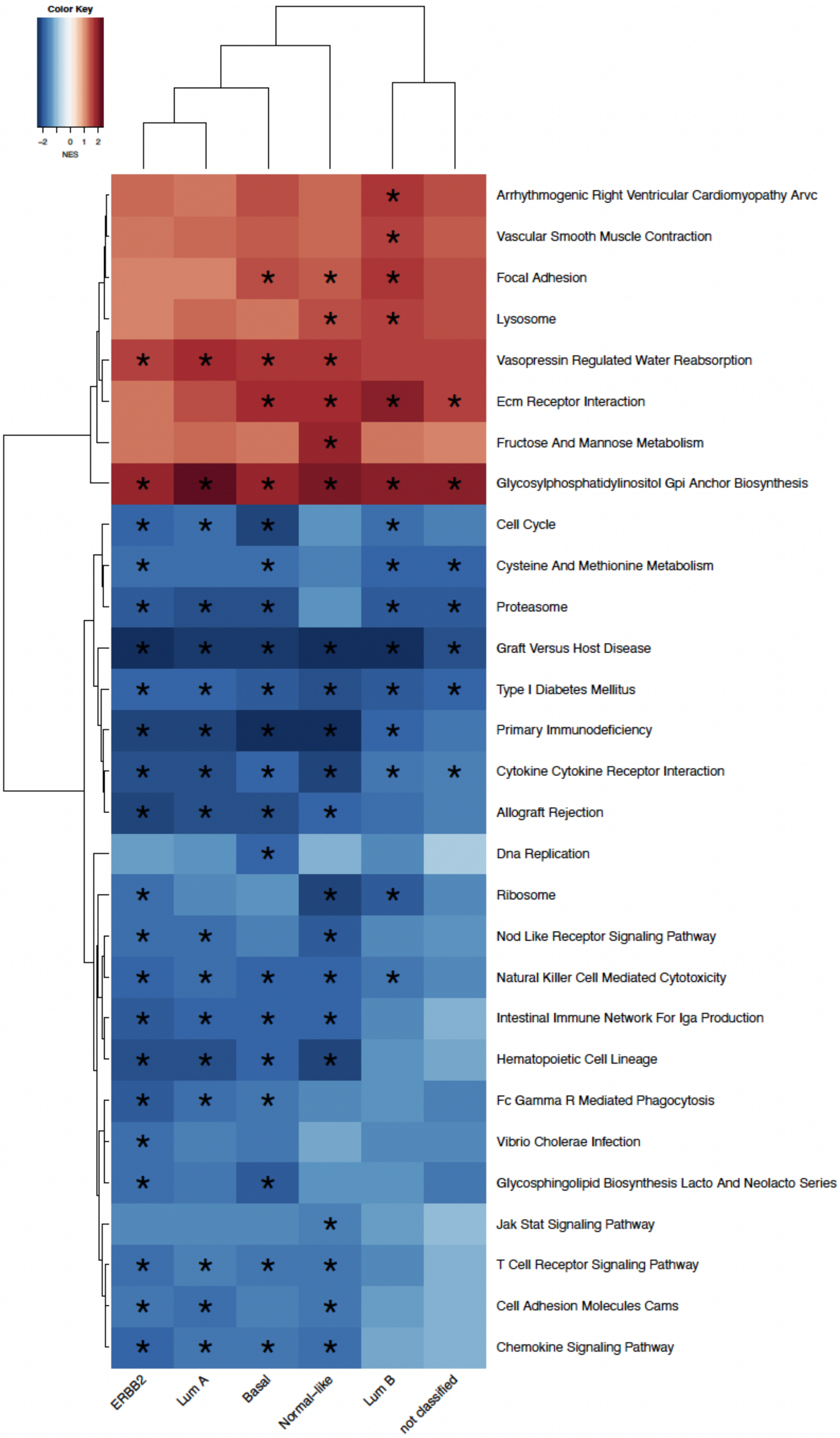

Figure S7: Pathways significantly (at FDR cutoff 0.05) correlated (positively or negatively) with miRNA expression in various breast cancer subtypes: heatmaps are colored by normalized enrichment scores (NES); Pathways positively correlated with miRNA expression are colored red and pathways negatively correlated with miRNA expression are colored blue.

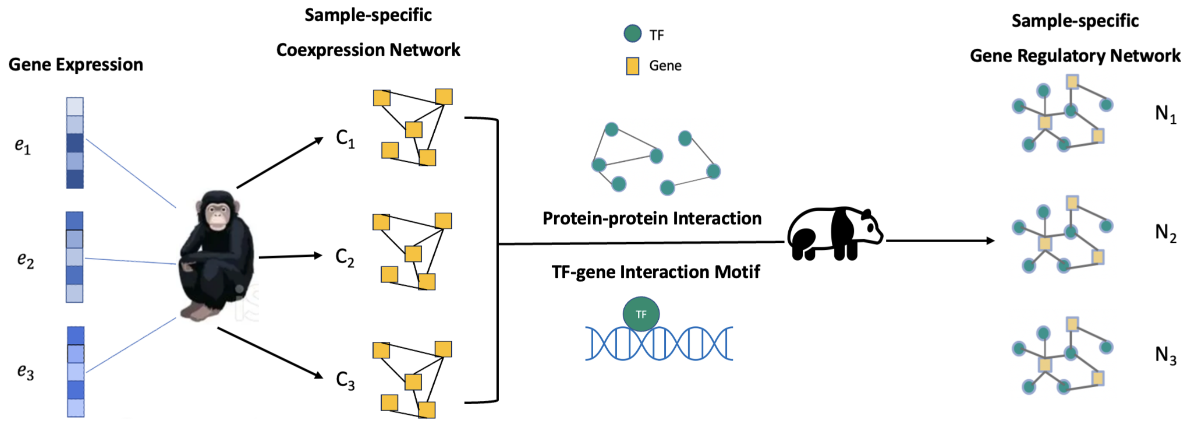

Figure S8: Schematic diagram of the BONOBO-PANDA pipeline: sample-specific co-expression matrices derived by BONOBO are used as input in the PANDA algorithm to derive sample-specific gene regulatory networks.

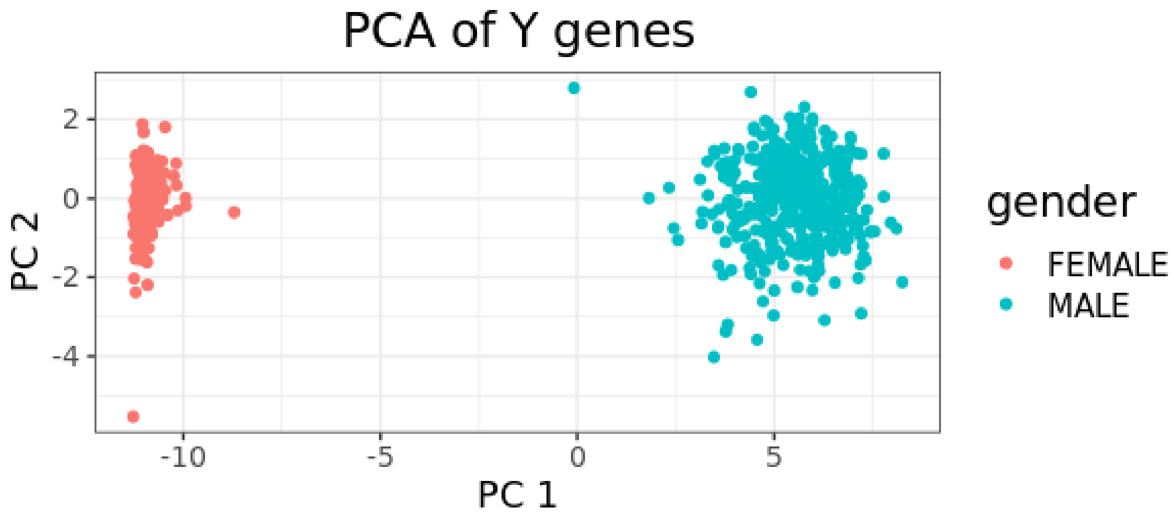

Figure S9: PCA of log transformed TPM normalized Y gene expression from GTEx thyroid samples: Self reported gender matches with chromosomal sex alignment in GTEx thyroid samples.

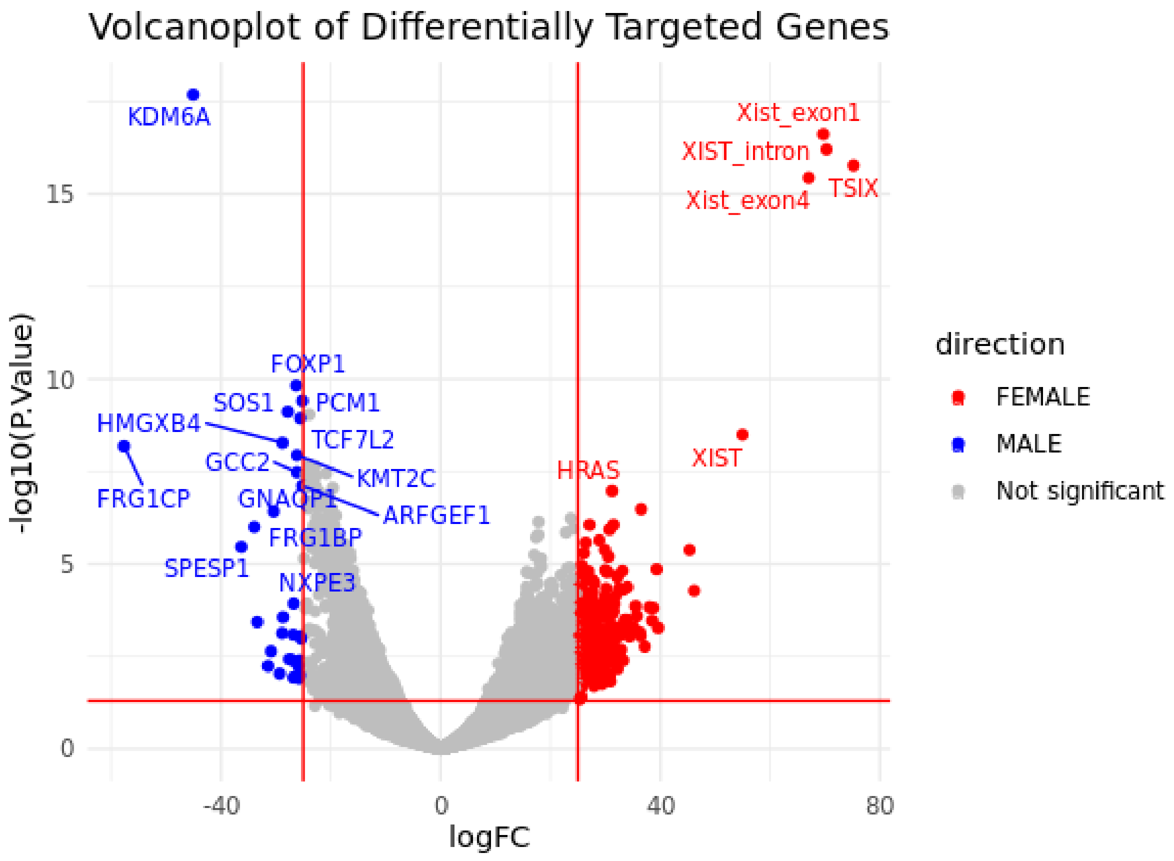

Figure S10: Genes most differentially regulated in males and females in GTEx thyroid samples: volcano plot from “limma” analysis.

#### S2 Supplementary Tables

Table S2.1: Mean squared error for BONOBO, LIONESS and SPCC on simulated data for varying number of genes. In paranthese we report the MSE scaled by the variance (last column) of the correlations in the true correlation matrix.

| **Number of Genes** | **BONOBO** | **LIONESS** | **SPCC** | **Variance** |
|---|---|---|---|---|
| 100 | 0.0151 (0.0933) | 0.1554 (0.9604) | 0.1732 (1.0705) | 0.1618 |
| 500 | 0.0169 (0.1673) | 0.1075 (1.0644) | 0.1125 (1.1139) | 0.1010 |
| 1000 | 0.0178 (0.2602) | 0.0812 (1.187) | 0.0835 (1.2208) | 0.0684 |
